# Impaired labyrinth formation prevents the establishment of the maternal-fetal interface in conditional Hand1-deficient mice

**DOI:** 10.1101/2020.09.02.280354

**Authors:** Jennifer A. Courtney, James Cnota, Helen Jones

**Affiliations:** Cincinnati Children’s Hospital Medical Center; University of Florida

## Abstract

**Introduction:** Congenital heart defects (CHD) affect approximately 1% of all live births, and often require complex surgeries at birth. Placental development and function is vital to ensure normal fetal development. We have previously demonstrated abnormal placental development and vascularization in human CHD placentas, and placental expression changes in genes important for heart development. Hand1 has roles in both heart and placental development and is implicated in CHDs including double right outlet, hypoplastic left heart syndrome, and septal defects; however, Hand1 involvement in placental vascularization and development is under-investigated. We utilized the Hand1^A126fs/+^ murine model to investigate Hand1 in placentation and vascularization.

**Methods:** Hand1^A126fs/+^ female mice were time-mated with Nkx2.5^cre^ (placenta- and heart-specific) males to produce either Nkx2.5^cre^;Hand1^+/+^ or Nkx2.5^cre^;Hand1^A126fs/+^ fetuses. Feto-placental units were harvested at timepoints from E8.5 to E14.5 for histological analysis; vascular assessment by immunohistochemistry for Hand1, CD-31, and CK-7; and angiogenesis by qPCR.

**Results:** Embryonic lethality occurs in Nkx2.5^cre^/Hand1^A126fs/+^ by E14.5 due to a failure of placental labyrinth formation and vascularization. Chorionic trophoblasts did not form, although trophoblast giant cell subtypes were present. Fetal vessels failed to develop properly and were significantly lower in the labyrinth by day E12.5. Placental growth factor levels were significantly increased, and Angiopoietin2 expression trended higher in Nkx2.5^cre/^Hand1 ^A126fs/+^ placental labyrinths compared to control littermates.

**Conclusion:** We demonstrate that Hand1 expression in placental chorion and trophoblast is necessary for proper patterning of the labyrinth and vascularization within the labyrinth. Multiple angiogenic factors known to be expressed in trophoblast were disrupted in Nkx2.5^cre/^Hand1 ^A126fs/+^ placental labyrinths compared to control littermates. Alterations in Hand1 expression represent a potential mechanism for abnormal placentation and early miscarriage in cases of CHD.

## Introduction

Successful establishment, development and function of the maternal-fetal interface ensures sufficient oxygen and nutrient transfer from maternal and fetal circulation throughout gestation, creating an optimal *in utero* environment for fetal development. Fetuses with congenital heart disease (CHD), the most common birth defect affecting ∼1% of all live births[1] demonstrate disrupted placental development and function [2, 3]. Despite decades of research with a primary focus on genetic etiology, the underlying cause of these defects remains unknown in the majority of cases[4]. Genome-wide associations studies (GWAS) undertaken by the Pediatric Cardiac Genomics Consortium (PCGC) have identified almost 400 genes thought to cause CHD[5, 6]. Population studies have recently demonstrated that pregnancies complicated by CHD carry a higher risk of developing pathologies associated with an abnormal placenta including growth disturbances[7-9], preeclampsia[10-13], preterm birth[14, 15], and stillbirth[16].

Modeling studies implicating genes responsible for abnormal heart development often overlook the involvement of extraembryonic tissues or circumvent it by using ‘cardiac-specific’ conditional knockout mouse models. However, this does not truly reflect the situation in cases of congenital heart defects where genetic perturbations would occur in all cells/tissues expressing that gene and many of the identified genes are expressed in other cell types in addition to those found in the heart. Importantly, in a recent study of murine models of heart development, 68% of 103 knockout models exhibiting embryonic lethality at or after mid-gestation had abnormal placental development[17], that had not previously been investigated, and there was a strong correlation between placental dysmorphology, angiogenesis and heart development.

Hand1 knockout mice were not examined in this study because of early lethality.

it has long been established that Hand1 is necessary for proper placentation and heart development in mouse. Placental and fetal development are arrested by gestational day 10 in Hand1-null fetuses because the placenta failed to form properly after implantation when trophoblasts fail to differentiate[18-21]. Engineered conditional knockout mouse models resolved many of the early lethality issues by isolating gene expression within the fetus. Despite this, some conditional Hand1 knockouts cause mid-gestation lethality, while others lead to viable offspring with mild phenotypes, depending on the cre-driver[22, 23], and these recent experiments focused solely on the fetus, despite the fact that some of the cre-drivers function in extra-embryonic cells of the yolk sac and placenta, which may explain the disparities in outcomes.

Modeling studies implicating genes responsible for abnormal heart development often overlook the involvement of extraembryonic tissues or circumvent it by using ‘cardiac-specific’ conditional knockout mouse models. However, this does not truly reflect the situation in cases of congenital heart defects where genetic perturbations would occur in all cells/tissues expressing that gene and many of the identified genes are expressed in other cell types in addition to those found in the heart. Importantly, in a recent study of murine models of heart development, 68% of 103 knockout models exhibiting embryonic lethality at or after mid-gestation had abnormal placental development[17], that had not previously been investigated, and there was a strong correlation between placental dysmorphology, angiogenesis and heart development.

Studies in the differentiation of human trophoblast stem cells syncytiotrophoblast demonstrate Hand1 gene expression, while isolated human placental cytotrophoblasts and villous tissue from deliveries at term had no Hand1 protein expression, suggesting that Hand1 is only required during early placental development[24, 25]. Similarly, human trophoblast cell models demonstrated variability in Hand1 expression, with BeWo and Jeg-3 cells expressing Hand1, while Jar and ED27 cells did not[26]. In the mouse, Hand1-null trophoblast stem cells failed to differentiate into trophoblast giant cells (TGC) and exhibited reduced invasion[27]; however, no studies were performed in the other labyrinthine trophoblast types.

In the current study, we sought to determine the placental contribution to embryonic lethality in Hand1-mutant mice under the Nkx2.5cre driver. Nkx2.5 is a well-known cardiac development gene, but is also required for yolk sac angiogenesis and is expressed in extraembryonic tissue[28, 29]. Previous experiments characterized the heart phenotype in this model, but did not investigate extraembryonic contributions to embryonic lethality[22].Critical to promoting organ development; driving fetal growth; and mitigating environmental exposures during pregnancy, the placenta has an important but understudied role in fetal CHD. The placenta is the vital organ during pregnancy which maintains the supply of oxygen and nutrients during fetal development, and we have previously demonstrated placental abnormalities at delivery in placentas of humans born with with complex congenital heart defects[30, 31]. Both the heart and placenta are vascular organs and develop concurrently[32], and shared pathways direct the development of both. Given the common developmental window and shared developmental pathways of the heart, placenta, and vasculature, we hypothesize that failure of proper placental development and and its reduced ability to properly nourish the developing fetus contributes to both human placental and cardiac anomalies. This study highlights the role of the placenta to properly form vascular beds in early pregnancy for fetal survival.

## Methods

### Cre Expression

All animal procedures were performed under protocols approved by the Institutional Animal Care and Use Committee of CCHMC. Nkx2.5^IREScre^ expression was verified in Ai14 tdTomato mice (Jackson 007914) at timepoints E8.5, E9.5, E10.5 and E12.5 for the Nkx2.5^IREScre^ driver mouse strain (Jackson 024637). Placentas and embryos were fixed in 4% paraformaldehyde (PFA), 2.5% polyvinylpyrrolidone (PVP), in phosphate-buffered solution (PBS) for 4 hours at room temperature on a rocker plate. Tissue was rinsed in PBS and placed in 30% sucrose until fully infused, then embedded in OCT and stored at −80C until ready to cut. Blocks were warmed to −20C and 7µm sections were mounted onto slides for immunofluorescence.

### Mice Mating and genotyping

To verify to efficacy of the Nkx2.5cre, we crossed homozygous Nkx2.5cre[29] males with homozygous tdTomato [33] females and collected fetoplacental units at E8.9, E9.5, E10.5, E11.0, and E12.5. Conditionally activated Hand1^A126FS/+^ [22] females were time mated with homozygous Nkx2.5^IREScre^ males to produce litters containing Nkx2.5^cre^;Hand1^A126FS/+^ and Nkx2.5^cre^;Hand1^+/+^ embryos.

Embryos and placentas were collected at E8.5, E9.5, E10.5, E12.5, and E14.5. Genotyping was performed on all feto-placental units within the litter by removing part of the yolk sac (through day E10.5) or clipping the tail of fetuses. Fetal tissue was digested in 72.75 µL Direct PCR lysis buffer and 2.25 µL Proteinase K for 4 hours at 56°C, vortexed, then denatured for 30 minutes at 85°C. See Table S1 for genotyping primers and cycling programs. To detect the frameshift mutation at the Hand1 locus, the Hand1 PCR product was digested with the restriction enzyme FauI (New England Biolabs R0651) using 1 ul of restriction enzyme with 1.5 ul of PCR product, 2 ul CutSmart buffer, and 20ul RNAse-free water with the standard protocol, then run on an agarose gel.

### RNA expression via reverse transcription quantitative polymerase chain reaction (qPCR)

Half placentas (E10.5 to E1.5) were flash frozen in liquid nitrogen and stored at −80C. Tissue was homogenized, and RNA was extracted using a Qiagen RNAEasy Minikit per protocol. RNA quantification and quality control were assessed using Nanodrop Spectrophotometer (Thermo Fisher). From each sample 1 µg of RNA was converted to cDNA using the Applied Biosystems high-capacity cDNA conversion kit per protocol. cDNA was stored at −20C. Expression levels of mouse Angpt1, Angpt2, VegfA and Plgf were assayed using 1/40^th^ of the cDNA template and 300nm/L of forward and reverse primer in a 25 µL SYBR Green PCR Master Mix reaction, using the Applied Biosystems StepOne Plus Real-Time PCR System. Expression was normalized using mouse Rps20. Relative quantification expression levels were calculated by comparative CT method using StepOne software v2.3.

### Immunohistochemistry and immunofluorescence

Implantation sites at E8.5 and E9.5, and embryos and half placentas at E10.5, E12,5, and E14.5 were fixed [4% paraformaldehyde (PFA), 2.5% polyvinylpyrrolidone (PVP), in phosphate-buffered solution (PBS)] for 4 hours at room temperature on a rocker plate. Tissue was rinsed in PBS, placed in 70% ethanol, processed and embedded in paraffin.

Hematoxylin and Eosin staining was performed on 5um sections of placental and fetal tissue, including full implantation sites. Briefly, sections were deparaffinized, rehydrated, placed in hematoxylin for 45 seconds, washed in running tap water, placed in 80% ethanol, dipped 3 times in eosin followed by dehydration, clearing and mounting in xylene-based mounting medium.

Immunohistochemistry was performed on 5um serial sections of mouse placenta and heart tissue. Sections were deparaffinized, rehydrated, and antigen retrieval was performed using Target Retrieval Solution (Dako). Sections were washed and endogenous peroxidase activity was blocked using 3% hydrogen peroxide. Afterward, sections were washed and nonspecific binding was blocked with serum corresponding to the secondary antibody host species followed by overnight incubation with the primary antibody. Sections were washed and incubated with appropriate biotinylated secondary antibodies (Vector Laboratories, Inc.). Antibody binding was detected using the avidin-biotin complex (ABC) kit followed by enzyme substrate DAB (Vector Laboratories, Inc). Sections were counterstained briefly in hematoxylin before dehydration, clearing and mounting. Protein expression was observed by light microscopy (Nikon, Melville, NY). All antibodies, concentrations, incubation times, and secondaries are detailed in Table S1.

Immunofluorescence was performed on 5um serial sections of mouse placental and heart tissue. Sections were deparaffinized, rehydrated, and antigen retrieval was performed using Target Retrieval Solution (Dako). Sections were washed and nonspecific binding was blocked with serum corresponding to the secondary antibody host species followed by incubation with the primary antibody. Sections were washed and incubated with appropriate fluorochrome-conjugated secondary antibodies. Sections were counterstained briefly with DAPI before being mounted with antifade mounting media. All antibodies, concentrations, incubation times, and secondaries are detailed in Table S1.

### Placental Fetal Vessel Counts

Placenta labyrinth fetal vessel counts were performed manually after identification of vessels by immunohistochemistry against CD-31 from 10 high-powered fields across the labyrinth on 2 5 umsections at least 50 um apart from each placenta (n number per genotype and dam here) using Nikon software.

### Statistical analysis and data presentation

Data are presented as mean ± SEM. N number refers to the number of litters. Statistical analysis was performed according to student’s t-test (Prism GraphPad). Level of significance (P-value) was defined as 0.05 or less.

## Results

### Nkx2.5cre is expressed in yolk sac, trophoblast cells, and cardiomyocytes

To verify the spatiotemporal expression of Nkx2.5cre, Nkx2.5cre males were crossed with tdTomato females and fetoplacental sections were immunostained with Hand1 antibody. Nkx2.5cre is expressed in trophoblasts at E8.5 and overlaps with Hand1 expression (Supplemental Figure 1A). Nkx2.5cre and Hand1 are co-expressed in yolk sac, labyrinth trophoblast progenitor cells and syncytiotrophoblast at E9.5 (Supplemental Figure 1B) and E10.5 (Supplemental Figure 1C). Sinusoidal giant cells do not express Nkx2.5cre at E10.5.

Nkx2.5cre is strongly expressed in heart at E8.5 and E9.5, and Hand1 is localized to cardiac ventricular tissue at E9.5 (Supplemental Figure 1D).

### No Nkx2.5cre;Hand1^A126fs/+^ fetuses were recovered at E14.5

Marked fetal demise occurred after E10.5 in Nkx2.5^cre^;Hand1^A126fs/+^ fetuses (Figure 1). While Nkx2.5^cre^;Hand1^+/+^ littermates showed consistent survival at all timepoints, noNkx2.5^cre^;Hand1^A126fs/+^ fetuses survived to E14.5. The expected ratio of Nkx2.5^cre^; Hand1^+/+^ to Nkx2.5^cre^;Hand1^A126fs/+^ was 50% each. Nkx2.5^cre^;Hand1^A126fs/+^ fetuses were overrepresented at E10.5 (55%); however, they showed a marked decrease at E12.5 (36%) and no viable fetuses were recovered at E14.5. Resorptions were not genotyped due to a lack of available fetal tissue. Normal resorption for wildtype c57/B6J mice is <10%.

**Figure 1.**
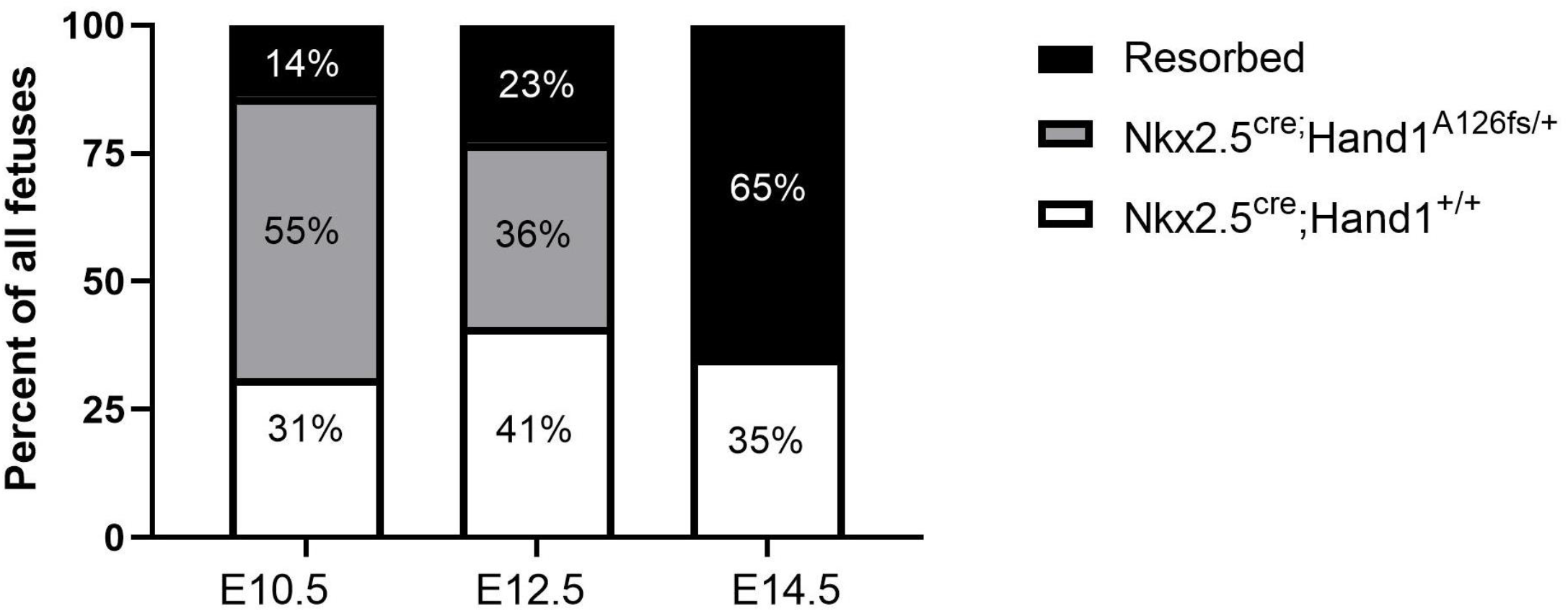
Nkx2.5^cre^;Hand1^A126FS/+^ show increased fetal demise. Graph shows percentages of live Nkx2.5^cre^;Hand1^+/+^ (white), Nkx2.5^cre^;Hand1^A126fs/+^ (grey), and resorbed (black) fetuses at E10.5, E12.5, and E14.5. No live Nkx2.5^cre^;Hand1^A126fs/+^ fetuses were retrieved at E14.5. N = 4-8 litters.

### Failure of labyrinthine formation in Nkx2.5^cre^;Hand1^A126fs/+^ placentas

By E9.5, labyrinthine morphogenesis is altered in Nkx2.5cre;Hand1A126fs placentas (Figure 2). In control littermate placentas (Figure 2A), the chorion exhibits invagination and folding with scattered Hand1-positive labyrinthine trophoblasts and a single layer of trophoblast giant cells separating labyrinth from decidua. Conditional knockout placentas (Figure 2B) have a disorganized layer of Hand1-positive trophoblast giant cells but lack Hand1-positive labyrinthine trophoblasts. Maternal blood spaces are present in both genotypes as indicated by positive cytokeratin-7 staining (not shown), but are dilated in the conditional knockouts compared to control littermates. In addition yolk sac morphology is altered in Nkx2.5^cre^;Hand1^A126fs/+^ fetuses at E9.5. The yolk sac is adjacent to the chorionic plate in the control littermates, but is separated from the trophoblast layer in Nkx2.5^cre^;Hand1^A126fs/+^ fetuses. Additionally, maternal blood spaces are dilated (expanded) in the knockouts compared to control littermates

**Figure 2.**
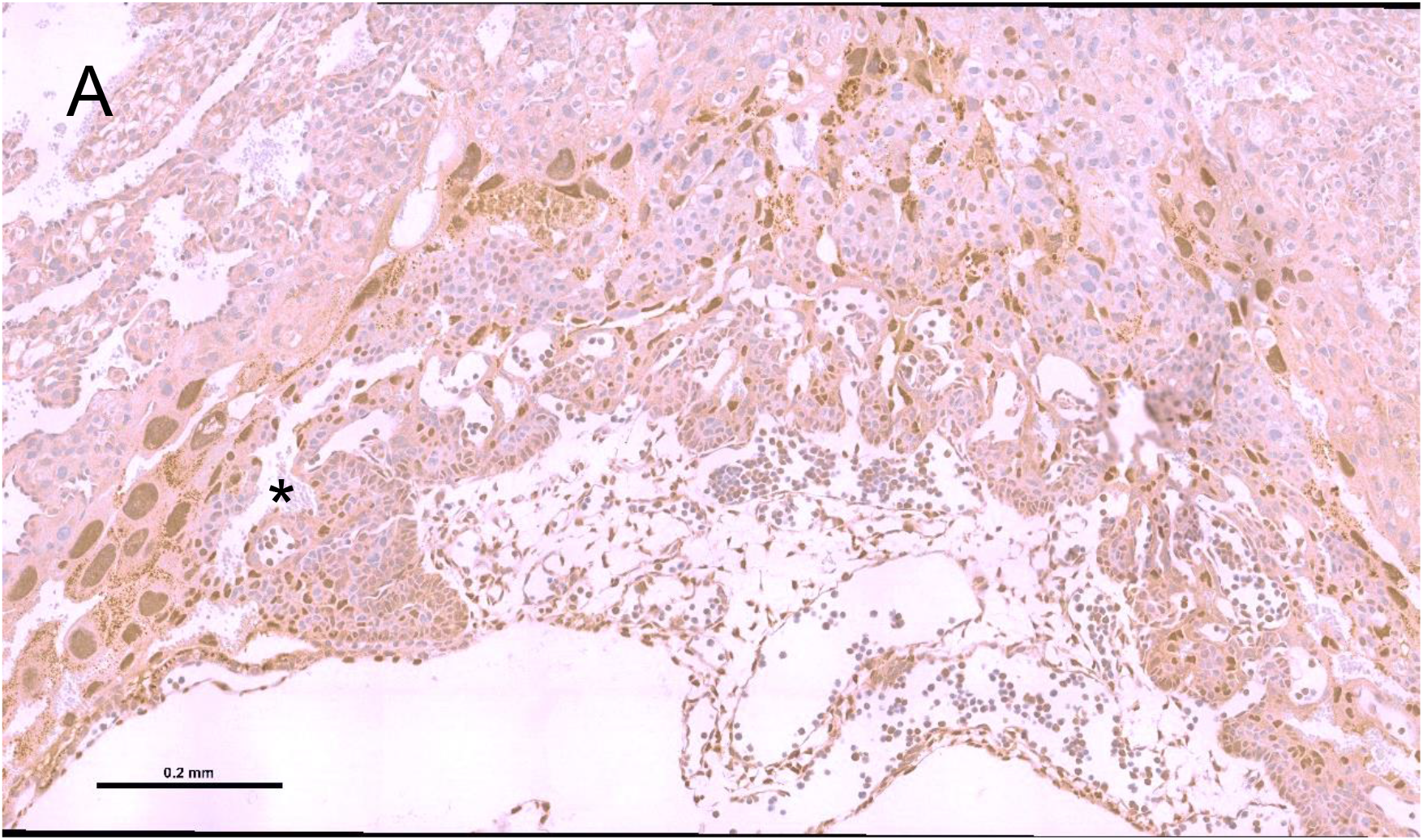
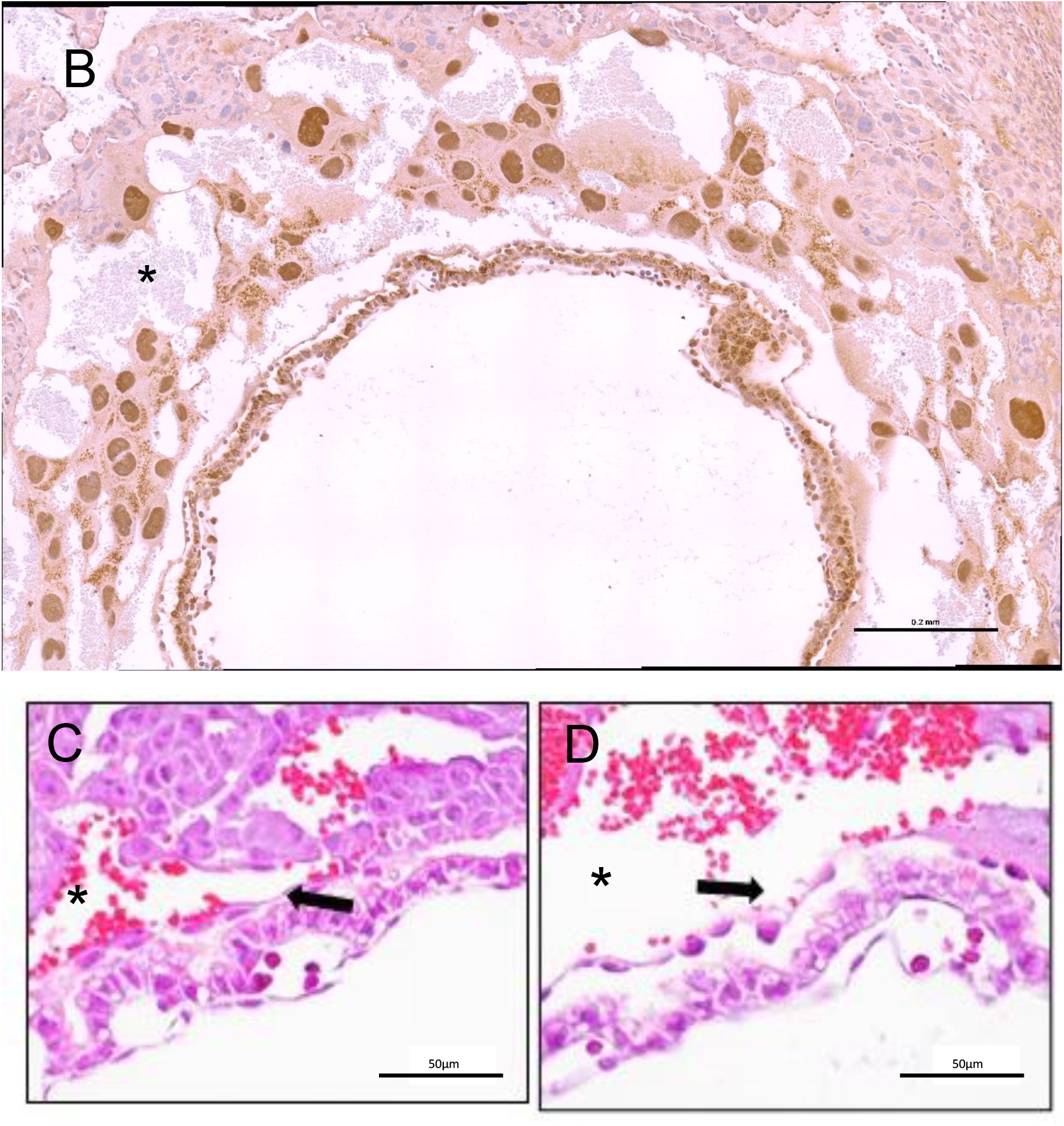
Labyrinthine morphogenesis is altered in Nkx2.5cre;Hand1A126fs/+ placentas at E9.5. (A) Control littermate placentas exhibit invagination of the chorion and development of an early labyrinth at E9.5. (B). In contrast, Nkx2.5cre;Hand1A126fs/+ placentas lack a chorionic plate and only have a disorganized layer of trophoblast giant cells surrounding the amniotic cavity. (C, D). Yolk sac morphology is altered in Nkx2.5^cre^;Hand1^A126fs/+^ fetuses at E9.5. (C). Normal littermates show adjacent chorion and yolk sac (arrow), while (D) Nkx2.5^cre^;Hand1^A126fs/+^ placentas lack a chorionic layer and yolk sacs are detached from the trophoblast layer separated by large maternal blood spaces (indicated by asterisks). (A,B) immunostained for Hand1. (C,D) stained with hematoxylin and eosin

### Failed labyrinth vascularization in Nkx2.5^cre^;Hand1^A126fs/+^ placentas

Nkx2.5^cre^;Hand1^A126fs/+^ fetuses show disruption in fetal labyrinthine vessel development (Figure 3). Fetal endothelium is not present in either the control littermates nor Nkx2.5^cre^; Hand1^A126fs/+^ labyrinths at E8.5 (not shown) and E9.5 (figure 3A,D), but angiogenesis is occurring by E10.5 only in the control labyrinths, as evidenced by the positive CD-31 staining for fetal endothelium (Figure 3B), whereas Nkx2.5^cre^;Hand1^A126fs/+^ placentas lack fetal labyrinthine blood vessels (Figure 3E). By E10.5, both syncytium and sinusoidal cytokeratin-7-positive trophoblast giant cells are present in the control labyrinth Figure 3B with clear delineation of fetal vasculature from maternal blood spaces, whereas the Nkx2.5^cre^;Hand1^A126fs/+^ labyrinths (Figure 3E) show disorganization of the trophoblasts and lack syncytium while retaining sinusoidal trophoblast giant cells. By E12.5, Nkx2.5^cre^;Hand1A^126fs/+^ placentas (Figure 3F) lack a fetal circulation, have very few remaining cytokeratin-7-positive trophoblasts, and the labyrinth layer has failed to develop compared to control (Figure 3C).

**Figure 3.**
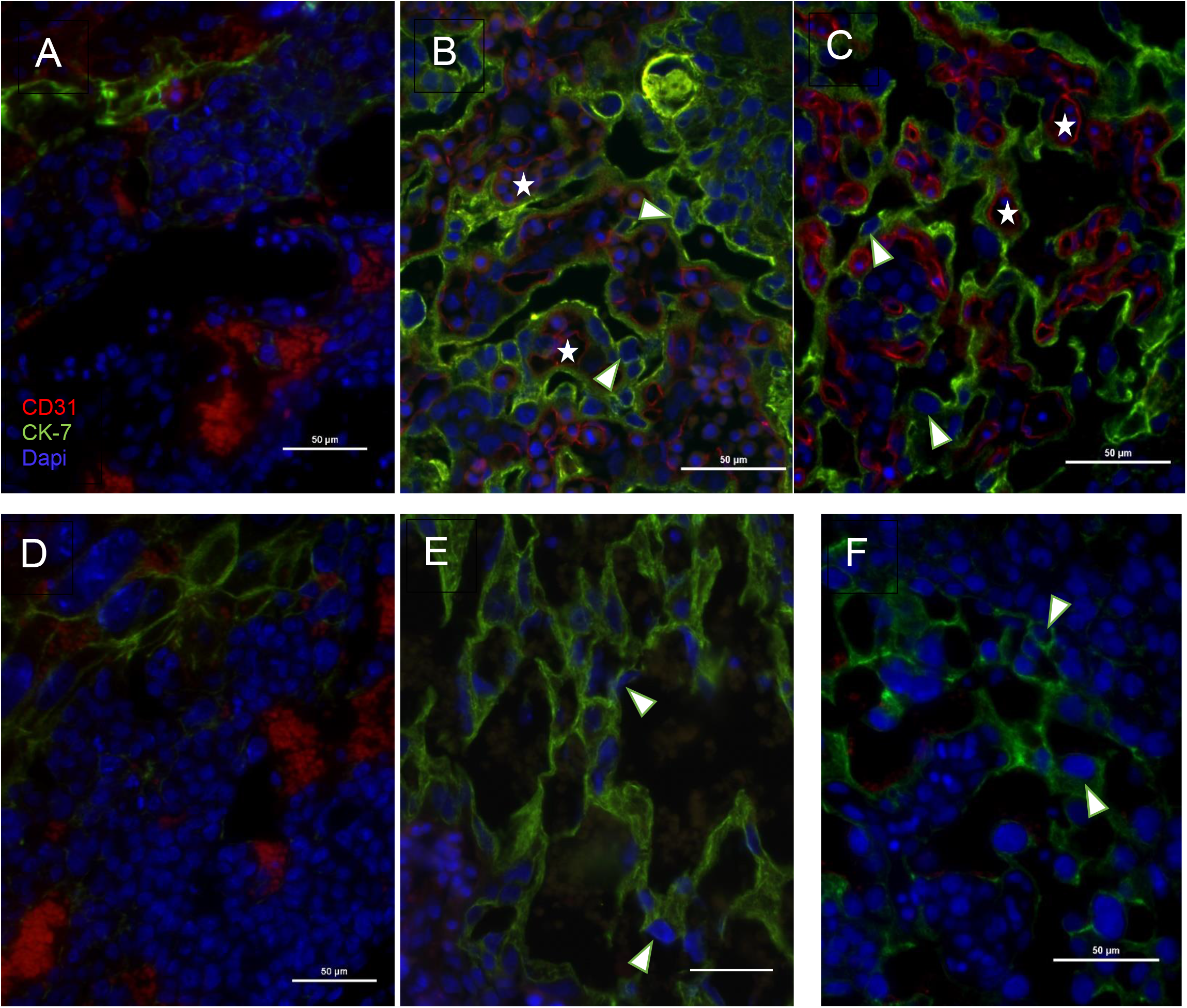
Nkx2.5^cre^;Hand1^+/+^ panels A-C, Nkx2.5^cre^;Hand1^A126fs/+^ panels D-F. Left column E9.5, middle E10.5, right E12.5. CD31-positive (stained red) fetal blood vessels are not present at E9.5 (A, D) in the chorionic plate for either genotype but both contain a layer of CK-7-positive (green) trophoblasts. Conditional Hand1 knockouts (E-F) show a disruption in syncytium with only sinusoidal trophoblast giant cells (arrowhead) lining the maternal blood spaces, while control littermates (B-C) have both a defined syncytium and labyrinthine fetal blood vessels.CD31-positive fetal endothelium (asterisk) is absent in conditional Hand1 knockouts at E10.5 and E12.5.. Red CD-31 (fetal endothelium), green Cytokeratin-7 (trophoblast), blue is DAPI.40X representative images.

### Fetal vessel counts were significantly reduced at E12.5 but not at E10.5 in Nkx2.5^cre^;Hand1^A126fs/+^ placental labyrinths

By gestation day 10.5, CD-31-positive fetal endothelium sparsely populated the labyrinth with a clear reduction in branching and the fetal vessels did not lay adjacent to maternal blood spaces in Nkx2.5^cre^;Hand1^A126fs/+^ placentas (Figure 4A,C). Fetal labyrinthine vessel counts were decreased in Nkx2.5cre;Hand1A126fs/+ placentas compared to control (13.6±1.4 vs. 7.4±3.4; p=0.088) with large variability in the conditional knockout labyrinths. By E12.5, vessel counts were significantly lower in Nkx2.5cre;Hand1A126f/+ labyrinths (31.8±3.4 vs. 2.6±1.2; p=0.001) compared to control.

**Figure 4.**
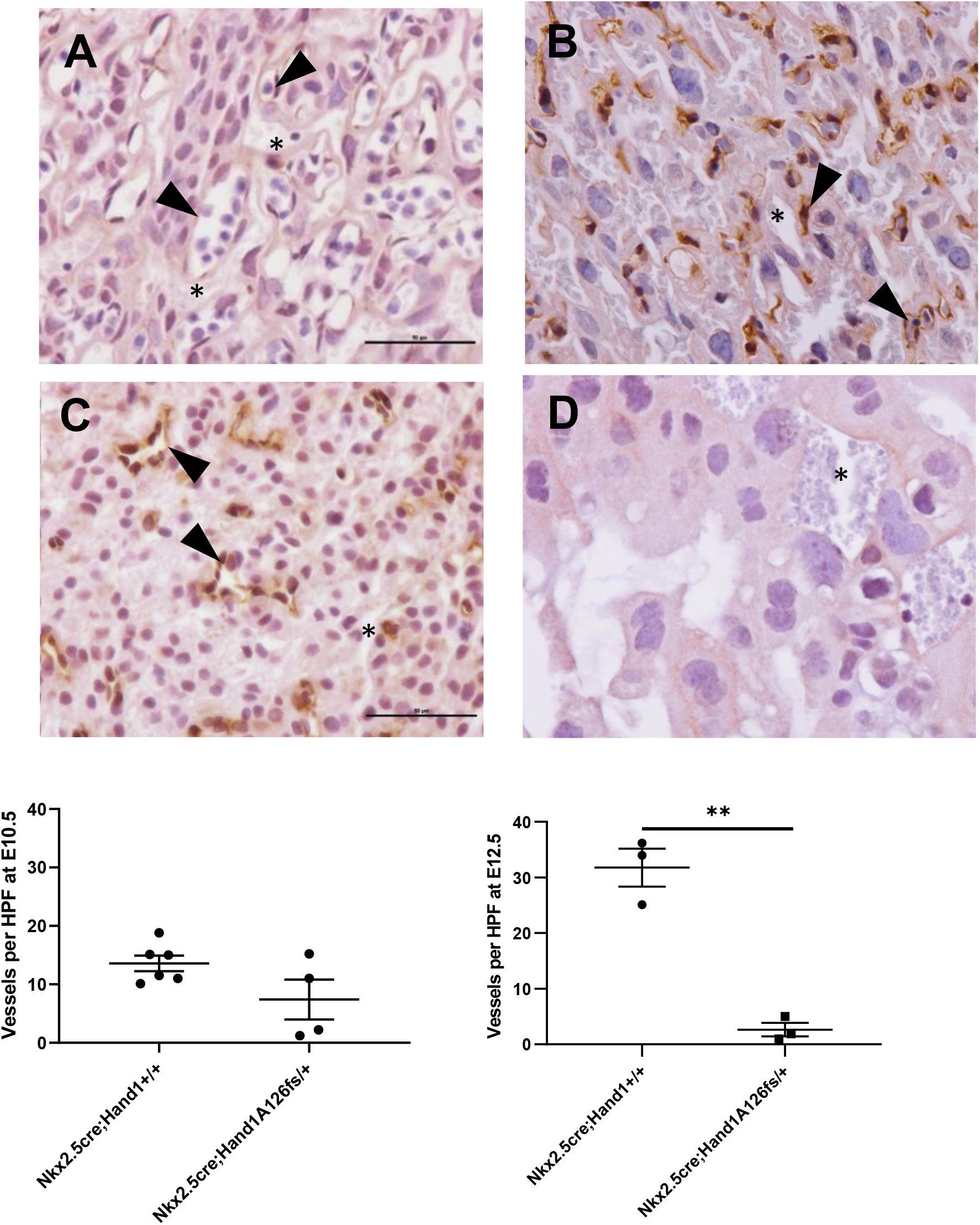
(B,D,F) Fetal blood vessel counts were significantly reduced in the Nkx2.5^cre^;Hand1^A126fs/+^ labyrinth at E12.5 compared to littermate controls (31.8±3.4 vs 2.6±1.2, p=0.001). (A,C,E) While not significantly different at E10.5, vessel counts trended downward at E10.5 (13.6±1.4 vs 7.4±3.4, p=0.088) with decreased branching and lack of contact with maternal blood spaces. Vessels were stained with CD-31 for fetal endothelium (brown). Arrowheads indicate fetal vessels, while asterisks mark maternal blood spaces. N= 4-6 litters, bars indicate mean ± SEM. ** p<0.01

### Gene expression

Plgf expression was significantly increased in isolated labyrinthine tissue from Nkx2.5^cre^; Hand1^A126fs/+^ compared to littermate controls at E10.5 (n=5 litters, p=0.02), Figure 5). In contrast there were no significant changes in gene expression for VegFa, VegFb, Angpt1, and Angpt2in Nkx2.5^cre^; Hand1^A126fs/+^ labyrinths at E10.5, although Angpt2 did trend higher (n=5 litters, p=0.08)… There were no significant sex differences in survival, labyrinth morphology, or fetal vessel density.

**Figure 5.**
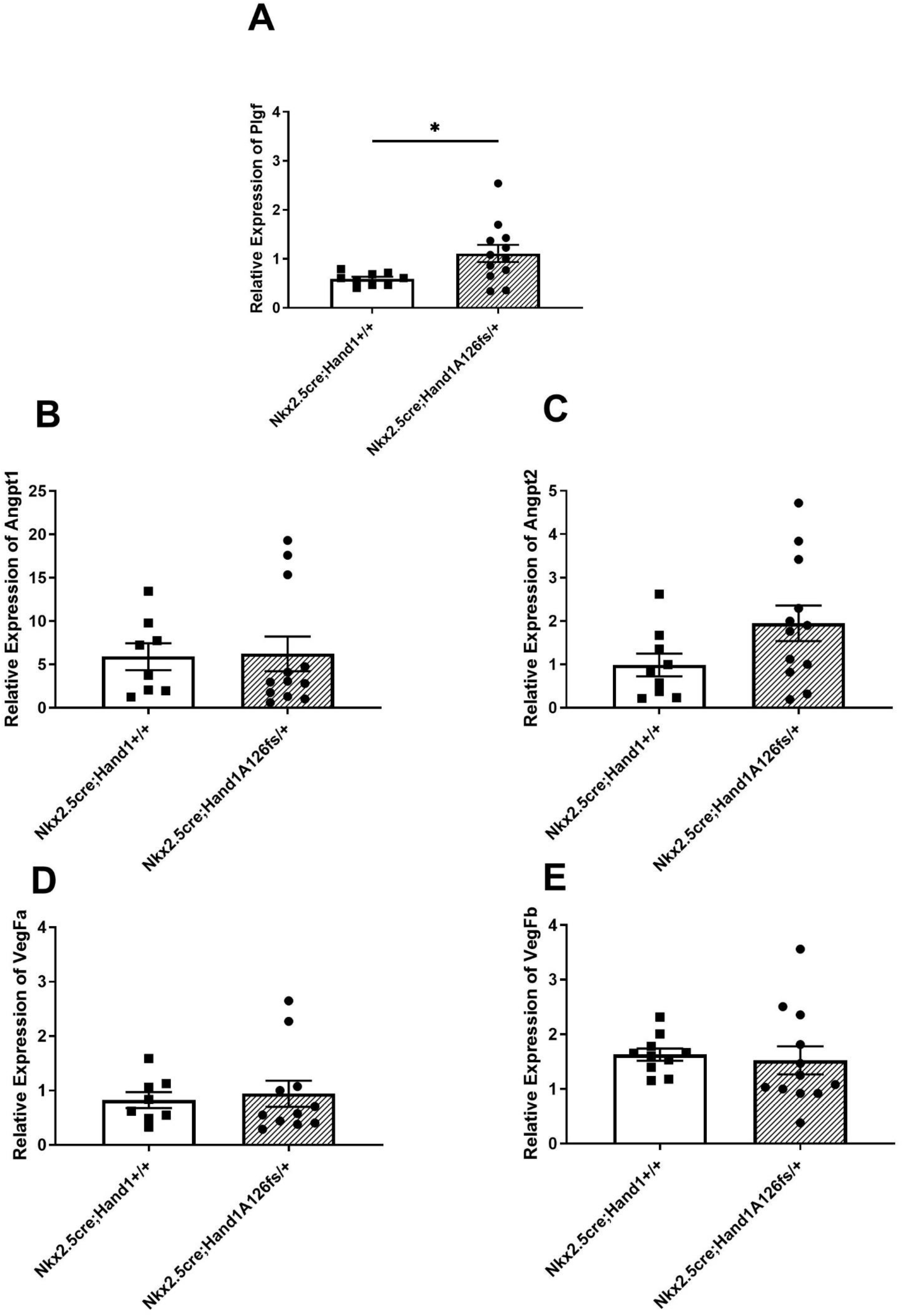
(A) Relative Plgf expression was significantly increased in the Nkx2.5^cre^; Hand1^A126fs/+^ labyrinths verses littermate controls. (B-E) Relative gene expression of VegFa, VegFb, Angpt1, and Angpt2 were not significantly different between Nkx2.5^cre^; Hand1^A126fs/+^ and Nkx2.5^cre^;Hand1^+/+^ placental labyrinthine tissue at E10.5, although Angpt2 trended higher in the conditional Han1 knockouts. Bars indicate mean ± SEM.

## Discussion

Targeted loss of Hand1 in chorionic and labyrinthine trophoblasts of the mouse placenta between gestation day 8.5-9.5 leads to abnormal formation of the placental labyrinth from both a trophoblast and endothelial perspective. Failure to form the appropriate layers of the maternal-fetal interface within the labyrinth (syncytium and vasculature) by E10/10.5 prevents the switch from histiotrophic to hemotrophic nutrition necessary for continued embryonic survival. While subsets of trophoblast cells of the labyrinth still received oxygen and nutrients carried in blood from maternal blood spaces, the failure to transfer nutrients across the syncytium and vascularize the labyrinth prevents them, and any diffused oxygen, from reaching the fetal circulation and results in embryonic lethality.

Our present study disrupted Hand1 expression under control of the Nkx2.5cre driver[29], with coexpression of the Nkx2.5 reporter and Hand1 limited to trophoblasts at gestation day 8.5 and expanded to include chorion by E9.5 and syncytium by E10.5. Sinusoidal giant cells and parietal cells did not express the Nkx2.5 reporter at E9.5 and beyond, but were Hand1-positive at these timepoints.

In contrast to studies in global Hand1 knockout mice[19, 26] and *in vitro* trophoblast stem cell differentiation to trophoblast giant cells [23], Hand1 expression was present in the trophoblast giant cells of Nkx2.5^cre^; Hand1^A126fs/+^ placentas at gestational day 9.5, suggesting that the timing of Hand1 disruption utilizing the NKx2.5-driven cre resulted in disruption to different populations of trophoblast cells allowing differentiation of trophoblast stem cells to TGCs (sinusoidal, canal, and parietal) for establishment of placentation while disrupting syncytialization and labyrinth formation.

In the current study the impaired development of the Nkx2.5^cre^;Hand1^A126fs/+^ placental labyrinth included disrupted vasculature in addition to abnormal trophoblast structure. In mice, the yolk sac undergoes *de novo* vascularization via the formation of blood islands[34], and disrupted fusion of the yolk sac with the chorionic plate that is seen at day 9.5 in our current study would deprive the developing labyrinth a source of endothelial cells. This results in a reduction in fetal labyrinthine vascularization, whereby oxygen and nutrients would not enter the fetal circulation, leading to embryonic lethality. Plgf, an important angiogenic factor in the placenta[35], was significantly upregulated in Nkx2.5^cre^;Hand1^A126fs/+^ placental labyrinth tissue, possibly in response to hypoxia signals from the placental tissue due to lower fetal vessel density. Additionally, Angpt2 – a competitor of Angpt1 to Tie2 receptors[36] – was increased, which would further destabilize any fetal endothelial vessels in the developing labyrinth. Interestingly, exposing wildtype dams to hypoxia at day E9.5[30] resulted in frequent fetal cardiac defects, highlighting the importance of sufficient placental oxygen uptake to heart development. Similarly, lower fetal oxygen uptake due to disrupted fetal placental angiogenesis in Nkx2.5^cre^;Hand1^A126fs/+^ fetuses may account for both the worsened cardiac defects previously demonstrated in that study compared to controls[22], and the resulting early embryonic lethality.

Previous studies demonstrate that ventricular Hand1 deletion in cardiomyocytes alone only leads to fetal survival to term with only mild cardiac phenotypes[22]. These studies support the hypothesis that abnormal placental development due to Hand1 disruption plays a primary role in the continuation of pregnancy and embryonic survival. The dual impact of a genetic disruption to the development of embryonic (heart) and extra-embryonic (placenta, yok sac) organs plus *in utero* environmental disruptions in oxygen and nutrient supply secondary to abnormal placenta development, may provide a mechanism that underlies early fetal loss. This may also underlie the high rate of miscarriage in humans associated with CHD directly or in prior/future pregnancy[37].

While mouse and human placentas have several differences, the cell types disrupted in this study have homologous cell types and functions in the human placenta[38]. The early fetal loss in this model correlates with first trimester miscarriage in human pregnancy and this targeted deletion created too severe of a phenotype to study impacts on the placenta seen in later pregnancy in cases of CHD[7-16]; therefore, other models will need to be developed. The role that the placenta may play in contributing to or exacerbating the development of CHD remains understudied, and many genes previously associated with CHD[4, 5] have not been adequately investigated in placental development or function. The DMDD mouse screen[17], and our study of placental gene expression in human CHD samples identifying multiple ‘heart-specific’ pathways from amongst the differentially expressed genes[31, 39], and underscore the importance of understanding the roles of developmental genes shared between placenta and heart.

By assessing placental development in the setting of a previously developed Hand1 mutation know to result in cardiac defects[22], we begin to explore the mechanisms that result in adverse pregnancy outcomes in the setting of fetal CHD. Failure of critical stages in placental development could lead to the known clinical observations of stillbirth, fetal growth restriction, and prematurity. As these outcomes occur in a variety of cardiac phenotypes and genetic mutations, further experiments are needed to understand the broad overlap of placental and cardiac development. Ultimately, such understanding may drive novel therapies to improve outcomes for children with CHD.

---

**Supplemental Figure 1.**
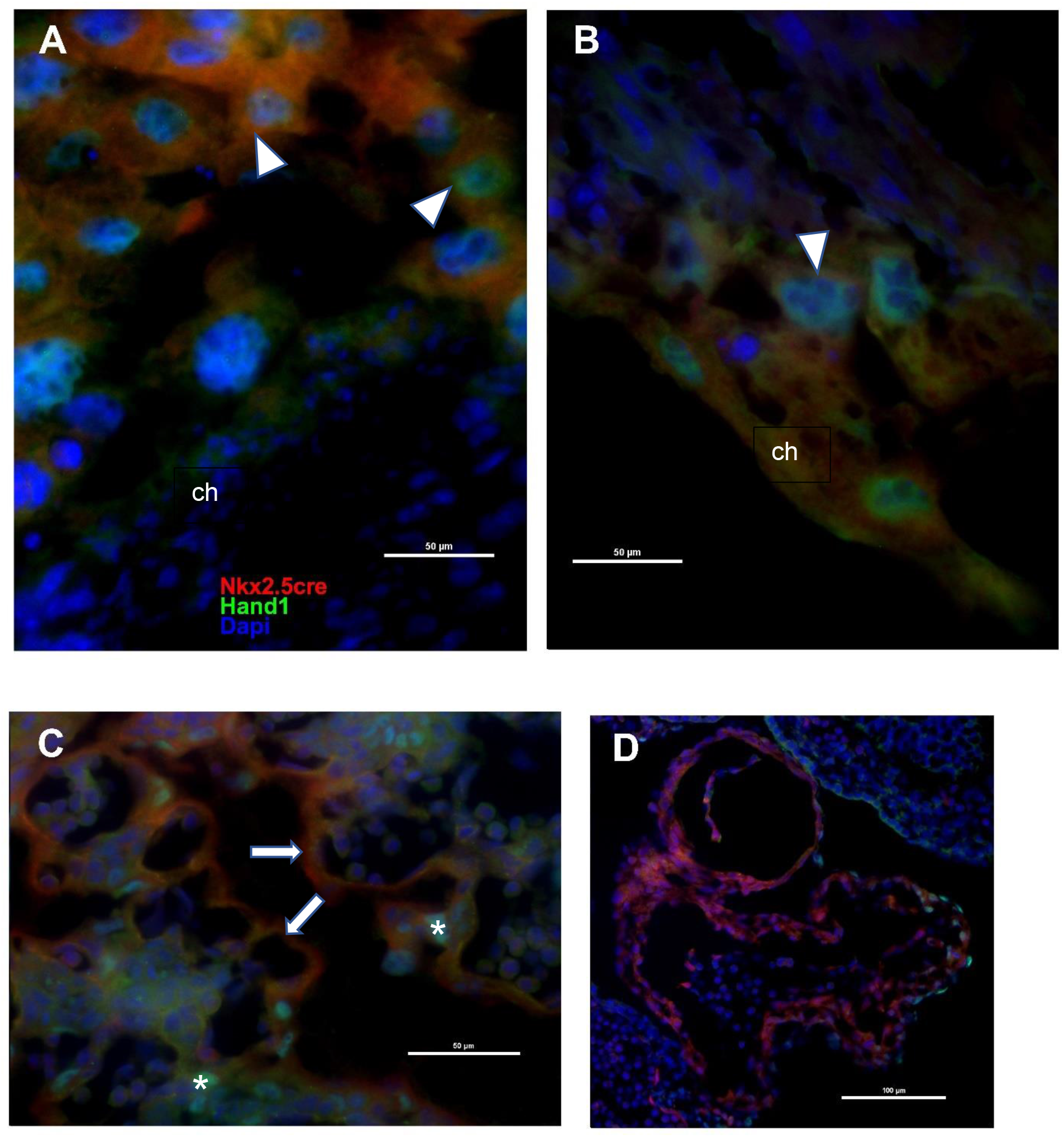
(A) Nkx2.5 (red) is expressed in trophoblast progenitor cells at E8.5, overlapping with Hand1 (green) protein expression, but not in chorion which does express Hand1. (B) At E9.5, Nkx2.5cre and Hand1 are coexpressed in both chorion and labyrinth trophoblast progenitor cells. (C) Hand1 and Nkx2.5cre overlap in syncytiotrophoblasts of the placental labyrinth and E10.5. Sinusoidal giant cells do not express Nkx2.5cre at E10.5. (D) Hand1 is coexpressed in a subset of ventricular cardiomyocytes at E9.5. Arrowhead is trophoblast progenitor cell. Arrow is syncytiotrophoblast. Asterisk is sinusoidal giant cell. ch = chorion

## References

1. van der Linde, D., et al., Birth prevalence of congenital heart disease worldwide: a systematic review and meta-analysis. J Am Coll Cardiol, 2011. 58(21): p. 2241–7.

2. Rychik, J., et al., Characterization of the Placenta in the Newborn with Congenital Heart Disease: Distinctions Based on Type of Cardiac Malformation. Pediatr Cardiol, 2018. 39(6): p. 1165–1171.

3. Burton, G.J., A.L. Fowden, and K.L. Thornburg, Placental Origins of Chronic Disease. Physiol Rev, 2016. 96(4): p. 1509–65.

4. Zaidi, S. and M. Brueckner, Genetics and Genomics of Congenital Heart Disease. Circ Res, 2017. 120(6): p. 923–940.

5. Jin, S.C., et al., Contribution of rare inherited and de novo variants in 2,871 congenital heart disease probands. Nat Genet, 2017. 49(11): p. 1593–1601.

6. Homsy, J., et al., De novo mutations in congenital heart disease with neurodevelopmental and other congenital anomalies. Science, 2015. 350(6265): p. 1262–6.

7. Cnota, J.F., et al., Somatic growth trajectory in the fetus with hypoplastic left heart syndrome. Pediatr Res, 2013. 74(3): p. 284–9.

8. Puri, K., et al., Fetal somatic growth trajectory differs by type of congenital heart disease. Pediatr Res, 2018. 83(3): p. 669–676.

9. Ruiz, A., et al., Placenta-related complications in women carrying a foetus with congenital heart disease. J Matern Fetal Neonatal Med, 2016. 29(20): p. 3271–5.

10. Auger, N., et al., Association Between Preeclampsia and Congenital Heart Defects. JAMA, 2015. 314(15): p. 1588–98.

11. Boyd, H.A., et al., Association Between Fetal Congenital Heart Defects and Maternal Risk of Hypertensive Disorders of Pregnancy in the Same Pregnancy and Across Pregnancies. Circulation, 2017. 136(1): p. 39–48.

12. Brodwall, K., et al., Possible Common Aetiology behind Maternal Preeclampsia and Congenital Heart Defects in the Child: a Cardiovascular Diseases in Norway Project Study. Paediatr Perinat Epidemiol, 2016. 30(1): p. 76–85.

13. Llurba, E., et al., Maternal and foetal angiogenic imbalance in congenital heart defects. Eur Heart J, 2014. 35(11): p. 701–7.

14. Laas, E., et al., Preterm birth and congenital heart defects: a population-based study. Pediatrics, 2012. 130(4): p. e829–37.

15. Tararbit, K., et al., Assessing the risk of preterm birth for newborns with congenital heart defects conceived following infertility treatments: a population-based study. Open Heart, 2018. 5(2): p. e000836.

16. Jorgensen, M., et al., Stillbirth: the heart of the matter. Am J Med Genet A, 2014. 164A(3): p. 691–9.

17. Perez-Garcia, V., et al., Placentation defects are highly prevalent in embryonic lethal mouse mutants. Nature, 2018. 555(7697): p. 463–468.

18. Cserjesi, P., et al., Expression of the novel basic helix-loop-helix gene eHAND in neural crest derivatives and extraembryonic membranes during mouse development. Dev Biol, 1995. 170(2): p. 664–78.

19. Riley, P., L. Anson-Cartwright, and J.C. Cross, The Hand1 bHLH transcription factor is essential for placentation and cardiac morphogenesis. Nat Genet, 1998. 18(3): p. 271–5.

20. Morikawa, Y. and P. Cserjesi, Extra-embryonic vasculature development is regulated by the transcription factor HAND1. Development, 2004. 131(9): p. 2195–204.

21. Firulli, A.B., et al., Heart and extra-embryonic mesodermal defects in mouse embryos lacking the bHLH transcription factor Hand1. Nat Genet, 1998. 18(3): p. 266–70.

22. Firulli, B.A., et al., The HAND1 frameshift A126FS mutation does not cause hypoplastic left heart syndrome in mice. Cardiovasc Res, 2017. 113(14): p. 1732–1742.

23. Firulli, B.A., et al., HAND1 loss-of-function within the embryonic myocardium reveals survivable congenital cardiac defects and adult heart failure. Cardiovasc Res, 2020. 116(3): p. 605–618.

24. Yabe, S., et al., Comparison of syncytiotrophoblast generated from human embryonic stem cells and from term placentas. Proc Natl Acad Sci U S A, 2016. 113(19): p. E2598–607.

25. Knofler, M., et al., Human Hand1 basic helix-loop-helix (bHLH) protein: extra-embryonic expression pattern, interaction partners and identification of its transcriptional repressor domains. Biochem J, 2002. 361(Pt 3): p. 641–51.

26. Knofler, M., et al., Molecular cloning of the human Hand1 gene/cDNA and its tissue-restricted expression in cytotrophoblastic cells and heart. Gene, 1998. 224(1-2): p. 77–86.

27. Hemberger, M., M. Hughes, and J.C. Cross, Trophoblast stem cells differentiate in vitro into invasive trophoblast giant cells. Dev Biol, 2004. 271(2): p. 362–71.

28. Tanaka, M., et al., Complex modular cis-acting elements regulate expression of the cardiac specifying homeobox gene Csx/Nkx2.5. Development, 1999. 126(7): p. 1439–50.

29. Stanley, E.G., et al., Efficient Cre-mediated deletion in cardiac progenitor cells conferred by a 3’UTR-ires-Cre allele of the homeobox gene Nkx2-5. Int J Dev Biol, 2002. 46(4): p. 431–9.

30. Jones, H.N., et al., Hypoplastic left heart syndrome is associated with structural and vascular placental abnormalities and leptin dysregulation. Placenta, 2015. 36(10): p. 1078–86.

31. Courtney, J., Troja, W., Owens, K., Brockway, H., Hinton, A., Hinton, R., Cnota, J., Jones, H., Abnormalities of placental development and function are associated with the different fetal growth patterns of hypoplastic left heart syndrome and transposition of the great arteries. Placenta, 2020. Accepted for Publication.

32. Burton, G.J. and E. Jauniaux, Development of the Human Placenta and Fetal Heart: Synergic or Independent? Front Physiol, 2018. 9: p. 373.

33. Risebro, C.A., et al., Hand1 regulates cardiomyocyte proliferation versus differentiation in the developing heart. Development, 2006. 133(22): p. 4595–606.

34. Forrai, A. and L. Robb, The hemangioblast--between blood and vessels. Cell Cycle, 2003. 2(2): p. 86–90.

35. Achen, M.G., et al., Placenta growth factor and vascular endothelial growth factor are coexpressed during early embryonic development. Growth Factors, 1997. 15(1): p. 69–80.

36. Yuan, H.T., et al., Angiopoietin 2 is a partial agonist/antagonist of Tie2 signaling in the endothelium. Mol Cell Biol, 2009. 29(8): p. 2011–22.

37. Koerten, M.A., et al., Frequency of Miscarriage/Stillbirth and Terminations of Pregnancy Among Women With Congenital Heart Disease in Germany, Hungary and Japan. Circ J, 2016. 80(8): p. 1846–51.

38. Georgiades, P., A.C. Ferguson-Smith, and G.J. Burton, Comparative developmental anatomy of the murine and human definitive placentae. Placenta, 2002. 23(1): p. 3–19.

39. Courtney, J.A., J.F. Cnota, and H.N. Jones, The Role of Abnormal Placentation in Congenital Heart Disease; Cause, Correlate, or Consequence? Frontiers in Physiology, 2018. 9(1045).

