## Supplemental Table 1 for "Impaired labyrinth formation prevents the establishment of the maternal-fetal interface in conditional Hand1-deficient mice"

Table S1.

| **Genotyping Primers** | **Forward** | **Reverse** |
|---|---|---|
| Hand1 275-448 | 5'-GCCCAAACGAAAAGGCTCAG-3' | 5'-AGCACGTCCATCAAGTAGGC-3' |
| Sry | 5'-AACAACTGGGCTTTGCACATTG-3' | 5'-GTTTATCAGGGTTTCTCTCTAGC-3' |
| Nifα | 5'-TGCTGTGTTCTGGTCAGTCAAG-3' | 5'-CAAAGCAAATCTCCATGCTCGG-3' |
| **qPCR Primers** | **Forward** | **Reverse** |
| Angiopoietin 1 | | |
| Angiopoietin 2 | | |
| VegFa | | |
| Plgf | | |
| **Antibodies for IHC and IF** | **Source** | **Concentration/Incubation** |
| Hand1 | R&D Systems AF3168 | 1:100 |
| CD-31 | R&D Systems AF3628 | 1:100 |
| CD-31 | Abcam ab28364 | 1:100 |
| Cytokeratin-7 | Abcam ab199718 | 1:800 |
